# Opportunistic neuronal fuelling

**DOI:** 10.64898/2026.06.08.730908

**Authors:** Y. Contreras-Baeza, F. Baeza-Lehnert, A. von Faber-Castell, C. Glück, A. San Martín, L. Ravotto, B. Weber, L. F. Barros

## Abstract

Neurons can utilize either glucose or lactate, yet fuel selection during periods of activity remains unresolved. By combining optical monitoring of cytosolic pyruvate and NADH with mathematical modelling, we systematically evaluated mitochondrial fuelling in cultured neurons and acute brain slices. Under substrate concentrations typical of resting brain tissue, neurons rely almost exclusively on glucose, with a minor contribution from pyruvate. However, when extracellular lactate rises to levels mimicking tissue activity, glycolysis is inhibited and lactate becomes a major substrate, irrespective of ongoing neuronal activity. Ultimately, neuronal fuel selection is dictated not by internal energy demand, but by extracellular lactate availability, which fluctuates with local glial metabolism and systemic states such as exercise. These findings redefine our understanding of short-term metabolic flexibility in the brain and underscore the significant yet understudied role of extracellular pyruvate.

## Introduction

Average ATP turnover in brain tissue is one order of magnitude faster than that of the rest of the body, with much of this energy invested in the recovery of ion gradients challenged by neurotransmission and action potentials. Studies with radiotracers, chemical analysis, respirometry, and other techniques converged on the view that the adult brain is fuelled by glucose, and that overall, the brain is almost fully aerobic, with more than 90% of ATP produced in mitochondria. Positioned between circulation and neurons are glial cells, which in response to neuronal cues take up glucose and convert it to lactate, an energy rich metabolite that sustains neurotransmission as effectively as glucose.

The factors that make neurons select lactate over glucose or vice versa are unclear. Because of limited spatiotemporal resolution, conventional approaches are not suited to address activity-dependent fuelling. Higher-resolution techniques based on fluorescence microscopy have been introduced, which can monitor metabolic parameters in complex settings with a resolution of seconds (Baeza-Lehnert et al., 2018; Díaz-García et al., 2017; Rangaraju et al., 2014). Collectively, these studies have shown that neuronal activity stimulates both glycolysis and mitochondrial OxPhos, but did not address the impact of the local lactate surge that accompanies brain activity (Barros et al., 2023; Bonvento & Bolaños, 2021; Raichle & Mintun, 2006) nor did they consider pyruvate, which is below NMRS resolution. Underscoring the role of lactate, a recent study found that pharmacological blockage of monocarboxylate transporters (MCTs) perturbed synchronized synaptic transmission only in the presence of added lactate (Söder et al., 2025).

Neuronal mitochondria cannot metabolize glucose directly, and the same limitation applies to lactate (Rauseo et al., 2026). Once in the cytosol, both substrates are funnelled into pyruvate and NADH, the two intermediate pools that support mitochondrial energization (Fig. 1). Pyruvate enters the organelle to be oxidized via the tricarboxylic acid cycle, while the high-energy electrons of cytosolic NADH are transferred to matrix NADH mostly via the malate–aspartate shuttle (MAS) and to a lesser extent via the glycerol phosphate shuttle. The question of glucose versus lactate was first approached experimentally under resting conditions by comparing rates of radiotracer uptake in pure neuronal cultures (Bouzier-Sore et al., 2003, 2006), revealing that both substrates may be metabolized in parallel. Taking advantage of the spatiotemporal resolution afforded by genetically-encoded sensors, we have here extended this analysis to determine the effect of electrical stimulation and extracellular monocarboxylates on fuel selection. Neurons were co-cultured with astrocytes, which promote their functional and metabolic differentiation (Barres et al., 1990; Hasel et al., 2017; Mamczur et al., 2015; Zhang et al., 2014). Real-time monitoring of metabolites was combined with mathematical modelling, and key observations were reproduced in acute tissue slices. Our main claim is that whether neurons consume glucose or lactate is determined by extracellular lactate, not by neuronal demand.

**Figure 1.**
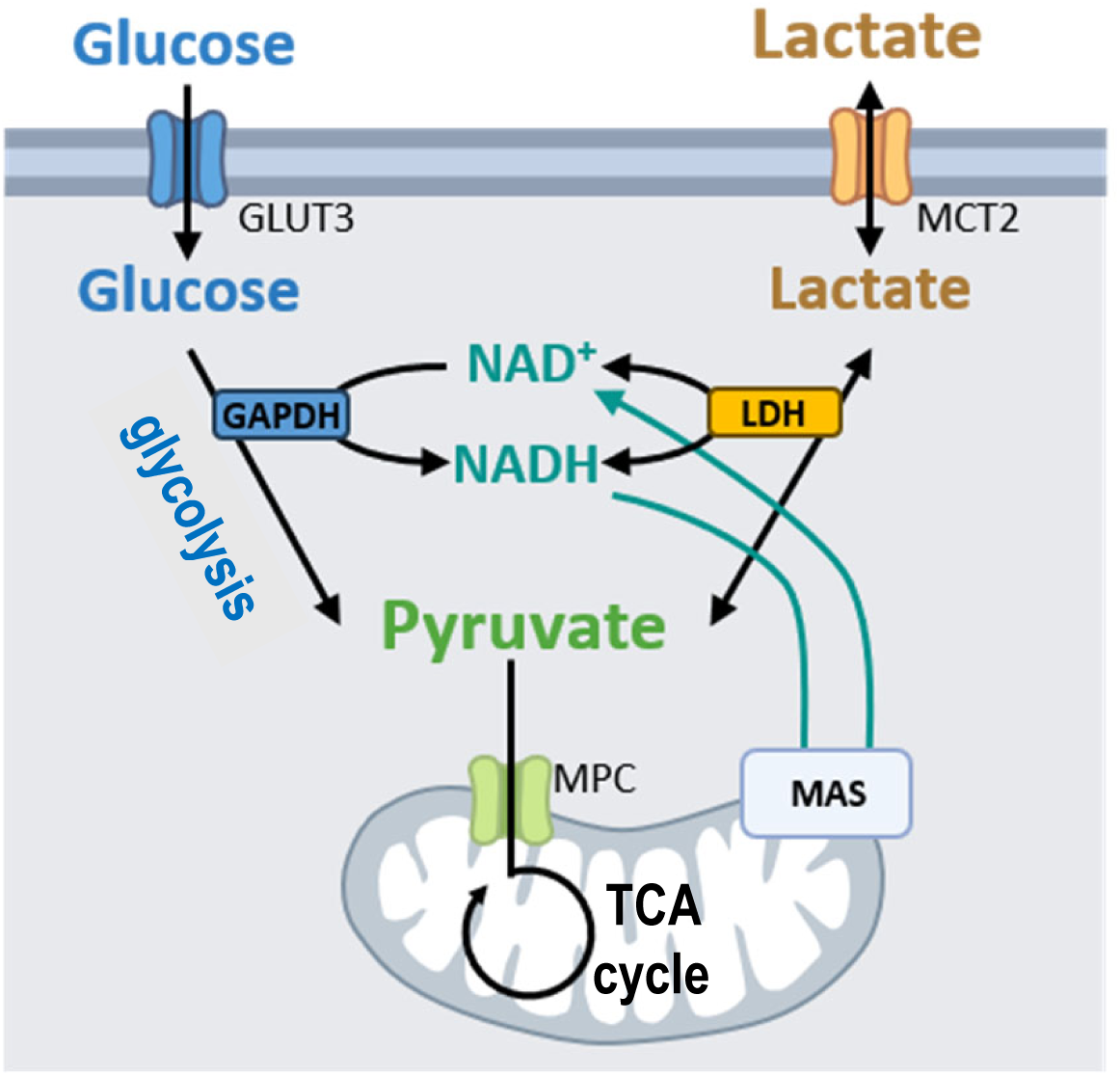
Neuronal fuelling. Neuronal mitochondria are energized by pyruvate and by NADH. Pyruvate is transported into mitochondria via the mitochondrial pyruvate carrier (MPC) to fuel the tricarboxylic acid (TCA) cycle. In parallel, reducing equivalents in NADH are transferred to the mitochondrial matrix via the malate–aspartate shuttle (MAS). The cytosolic pools of pyruvate and NADH are in turn sustained by glucose and by lactate. Glucose enters neurons via GLUT3 (and GLUT1) and is irreversibly metabolized to pyruvate by glycolysis. Lactate enters via MCT2 (and MCT1) and is reversibly metabolized to pyruvate by LDH. Glycolysis and LDH are redox-coupled at GAPDH reaction.

## Results

Unlike other cell types that process fatty acids and amino acids, neurons are solely fuelled by glucose and lactate. Most neuronal ATP is generated in mitochondria, and before fuelling the organelle, both substrates are transformed in the cytosol into pyruvate and NADH (Fig. 1). In view of this mandatory convergence, we posited that the question of neuronal fuelling could be approached by monitoring the composition of these two pools.

### MCT2 is near thermodynamic equilibrium

Experimental buffers for *in vitro* study of brain cells normally include glucose, sometimes lactate, but seldom pyruvate (Baeza-Lehnert et al., 2018; Díaz-García et al., 2017; Rangaraju et al., 2019). However, brain tissue has about 100 µM pyruvate (Conn & Steele, 1982; Hawkins et al., 1973), which is a better MCT1 & 2 substrate than lactate (Halestrap, 2012). Hence, we studied the influence of extracellular pyruvate on intracellular pyruvate. As shown in Figure 2A, the cytosolic pyruvate pool, measured with the FRET sensor Pyronic (San Martín et al., 2014), increased significantly when physiological pyruvate was added on top of physiological glucose and lactate.

**Figure 2.**
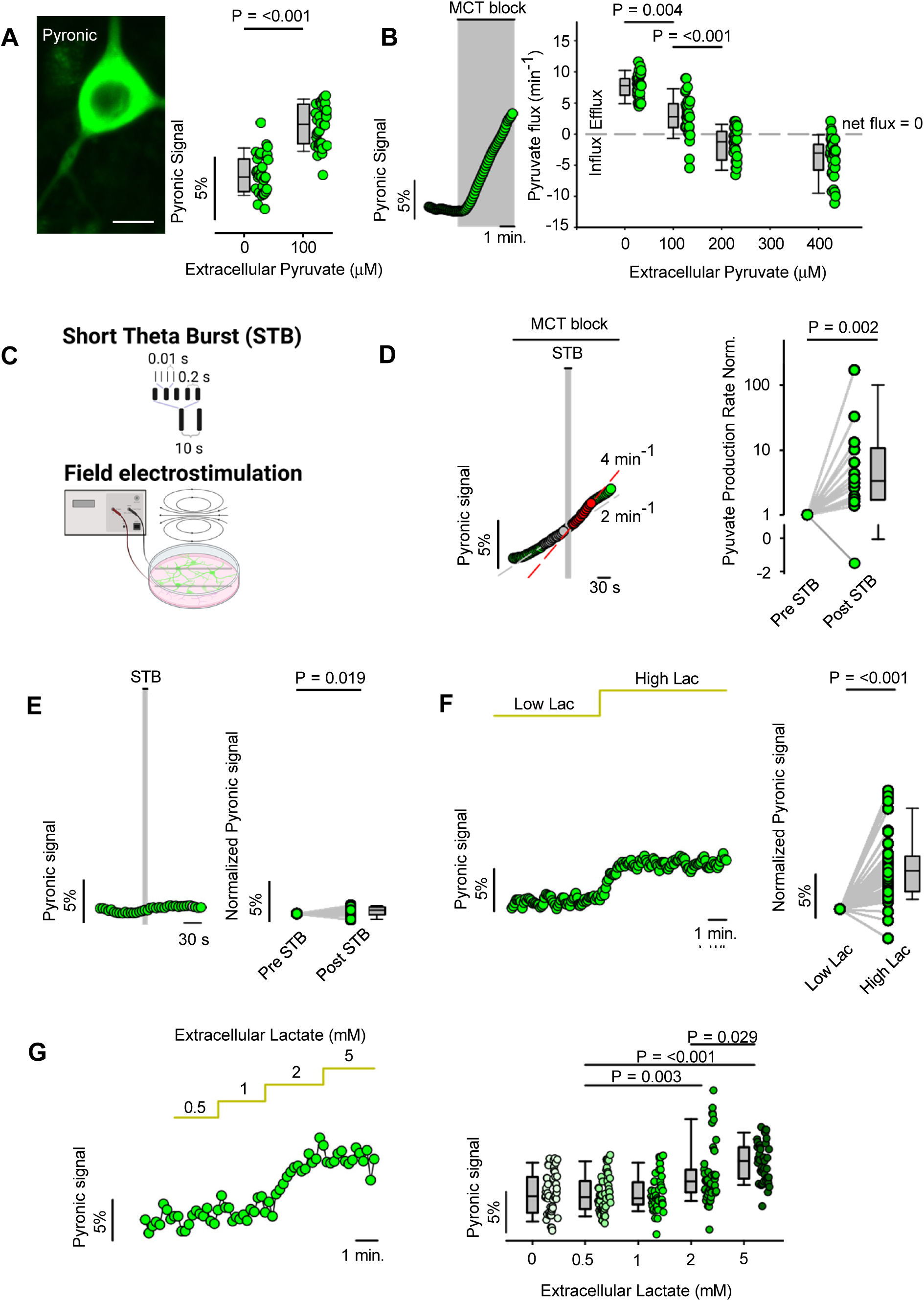
Neuronal pyruvate dynamics in response to electrical activity and extracellular pyruvate and lactate. **(A)** Impact of extracellular pyruvate on cytosolic pyruvate. Left panel: fluorescence image of a neuron expressing the FRET pyruvate sensor Pyronic. Bar = 10 µm. Right panel: summary of 5 experiments (31 cells) at zero pyruvate, and 6 experiments (34 cells) at 0.1 mM pyruvate. Perfusion solutions contained 2 mM glucose and 1 mM lactate. **(B)** Effect of extracellular pyruvate on net pyruvate flux through the plasma membrane, determined by exposing cells to the MCT1/2 blocker AR-C155858 (1 μM). Left panel: representative trace from a single neuron imaged in the absence of pyruvate. Right panel: summary of experiments performed at increasing extracellular pyruvate concentrations: 0 mM (n = 31 cells/5 experiments), 0.1 mM (n = 34 cells/6 experiments); 0.2 mM (n = 27 cells/4 experiments) and 0.4 mM (n = 37 cells/7 experiments). The dashed line indicates absence of net flux. Perfusion solutions contained 2 mM glucose and 1 mM lactate. **(C)** Schematic of field stimulation of hippocampal cultures using a short theta burst (STB) protocol. The STB is composed of two trains, separated by 10 s. Each train lasts 1 s and is composed of 20 square pulses of 1 ms duration, organized in 5 groups of 4 pulses, delivered at 0.2 s intervals. Individual pulses are 0.01 s apart. **(D)** Effect of STB on net pyruvate flux in cells exposed to the MCT blocker AR-C155858 (1 μM). Left panel: representative trace from a single neuron. Right panel: summary of slopes (n = 14 cells/7 experiments). The perfusion solution contained 2 mM glucose, 1 mM lactate and 0.1 mM pyruvate. **(E)** Effect of STB on cytosolic pyruvate level. Left panel: representative trace from a single neuron. Right panel: summary (n = 17 cells/ 12 experiments). The perfusion solution contained 2 mM glucose, 1 mM lactate and 0.1 mM pyruvate. **(F)** Cytosolic pyruvate was measured throughout the transition from low (0.5 mM) to high (5 mM) extracellular lactate. Left panel: representative trace from a single neuron. Right panel: summary of data (n = 48 cells/ 12 experiments). Perfusion solutions contained 2 mM glucose and 0.1 mM pyruvate. **(G)** Cytosolic pyruvate measured at increasing extracellular lactate concentrations (0 - 5 mM). Left panel: representative trace from a single neuron. Right panel: summary (n = 47 cells/ 5 experiments). The perfusion solution contained 2 mM glucose and 0.1 mM pyruvate.

MCT1 and MCT2 are blocked by AR-C155858 (Ovens et al., 2010), a useful tool for inhibitor-stop assessment of monocarboxylate fluxes across biological membranes. Because the cytosolic pools of pyruvate and lactate are tightly coupled by near-equilibrium LDH (Glancy et al., 2021; Gray et al., 2014; Krebs & Bellamy, 1960; Rogatzki et al., 2015), inhibitor-induced accumulation and depletion respectively reflect net monocarboxylate export and import (San Martín et al., 2014; Rauseo et al., 2026). In cultures supplied with glucose and lactate, without pyruvate, AR-C155858 induced cytosolic pyruvate accumulation, the rate of which decreased in the presence of physiological pyruvate and reverted to intracellular depletion at supraphysiological pyruvate levels (Fig. 2B). Thus, under physiological concentrations of glucose, lactate and pyruvate, cultured neurons lie at the tipping point between net monocarboxylate import and export. This finding might help to explain why previous studies, which omitted pyruvate and sometimes also lactate, arrived at contrasting conclusions regarding neuronal lactate usage.

### The pyruvate pool is sensitive to extracellular lactate, not to neurotransmission

To investigate the metabolic effects of neurotransmission, neurons were field stimulated using a short theta burst (STB), a protocol that mimics moderate hippocampal activity. The STB consists of two trains of impulses separated by 10 seconds (Figure 2C). Each train lasts for 1 second and is composed of twenty pulses of 1 ms duration, distributed into five groups of four (Albensi et al., 2007). The STB evokes a Ca^2+^ transient (see below) followed by a slower Na^+^ rise that correlates well with ATP demand (Baeza-Lehnert et al., 2018). In physiological pyruvate, the STB caused an increase in the rate of pyruvate accumulation (Fig. 2D), of a similar magnitude to that observed in the absence of pyruvate (Baeza-Lehnert et al., 2018). Also, in line with our previous study, the STB did not affect the size of the pyruvate pool, meaning that the rise in the export of monocarboxylates elicited by the stimulation is exactly matched by a rise in monocarboxylate production (Fig. 2E). These observations indicate that extracellular pyruvate, while influencing the size of the neuronal pyruvate pool, does not affect the metabolic response to neurotransmission.

Neural activity in brain tissue is accompanied by a quick increase in extracellular lactate. To investigate the possible impact of this factor on the neuronal pyruvate pool, cultures were switched from 0.5 mM to 5mM lactate. As shown in Figures 2F and 2G, this imposed rise in extracellular lactate had a concentration-dependent effect on neuronal pyruvate.

### The NADH pool is sensitive to extracellular lactate, not to neurotransmission

The second pathway of mitochondrial energization in neurons is the shuttling of electrons via the cytosolic NADH pool (Fig. 1), which can be monitored with Peredox (Choi et al., 2012). We asked whether this pool reacted to activity and extracellular lactate. No changes were detected in response to STB stimulation. In contrast, a significant increase in NADH followed the transition from low to high lactate (Figs. 3A-B). Neurons were then imaged at varying degrees of stimulation using Ca^2+^ as a proxy of metabolic demand (Fig. 3C). As shown in Figure 3D, cytosolic NADH levels did not respond to moderate stimulation, only increasing in response to tetanic stimulation and glutamate exposure. The lack of response was unexpected because a previous study using the same sensor reported activity-dependent NADH transients in neurons (Díaz-García et al., 2017). To investigate this discrepancy, we repeated the experiment with glucose as the only substrate, and at 34 - 35 °C, approaching the conditions used in the previous study. Nevertheless, physiological electrical stimulation had no detectable effect on NADH under either low- or high-lactate (Fig 3E). We do not have an explanation for the different behaviour, but there are multiple differences between the two studies, e.g. pyramidal cells versus granule cells, culture versus acute slices, and the presence of lactate and pyruvate in the perfusate.

**Figure 3.**
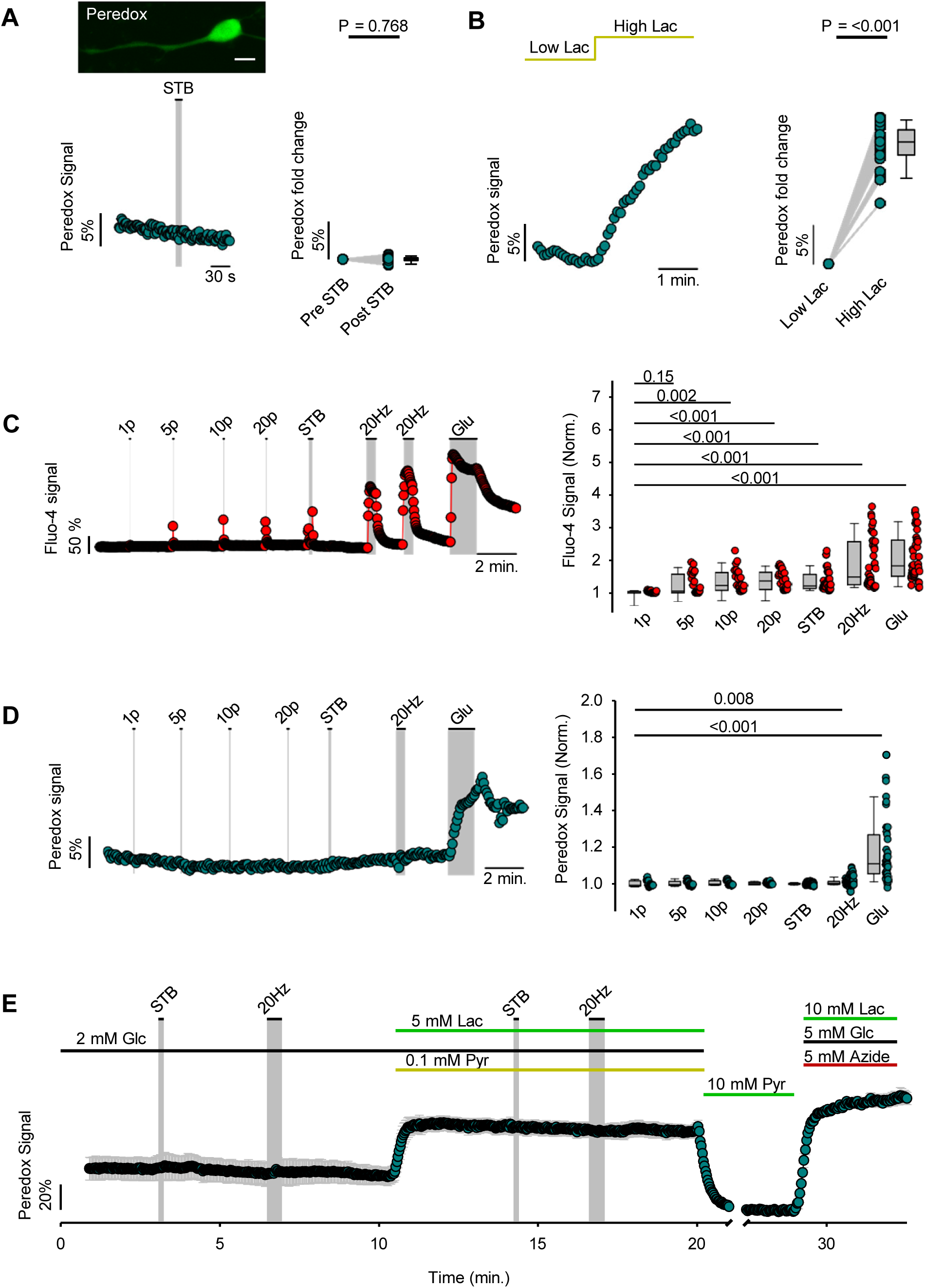
Neuronal NADH dynamics in response to electrical activity and extracellular lactate. **(A)** Top panel: fluorescence image of a neuron expressing the NADH/NAD^+^ sensor Peredox. Bar = 10 µm. Bottom panel: effect of STB on cytosolic NADH level. Representative trace from a single neuron. Right panel: summary of data (n = 24 cells/3 experiments). The perfusion solution contained 2 mM glucose, 1 mM lactate and 0.1 mM pyruvate. **(B)** Cytosolic NADH measured through the transition from low (0.5 mM) to high (5 mM) extracellular lactate. Left panel: representative trace from a single neuron. Right panel: summary of data (n = 24 cells/3 experiments). The perfusion solution contained 2 mM glucose and 0.1 mM pyruvate. **(C)** The effect of increasing number of electrical pulses on cytosolic Ca^2+^ was assessed using Fluo-4, from 1, 5, 10, 20, 40 (STB), 600 (20 Hz), and finishing with 50 µM glutamate. Left panel: representative trace from a single neuron. Right panel: summary of data (n = 22 cells/3 experiments for 1 - 20 pulses; n = 37 cells/6 experiments for 40 pulses; n = 40 cells/7 experiments for 20 Hz and 50 µM glutamate). The perfusion solution contained 2 mM glucose, 0.5 mM lactate and 0.1 mM pyruvate. **(D)** The effect of increasing number of electrical pulses on cytosolic NADH was assessed using Peredox, from 1, 5, 10, 20, 40 (STB), 600 (20 Hz), and finishing with 50 µM glutamate. Left panel: representative trace from a single neuron. Right panel: summary of data (n = 17 cells/ 3 experiments for 1 - 20 pulses, 125 cells/ 26 experiments for 40 pulses and 20 Hz and 39 cells/ 9 experiments for 50 µM Glutamate). The perfusion solution contained 2 mM glucose, 0.5 mM lactate and 0.1 mM pyruvate. **(E)** Effect of electrical stimulation and extracellular substrates on cytosolic NADH (Peredox) at 34 - 35°C. Cells were subjected to an STB and tetanic stimulation (20 Hz for 30 s) in the presence of glucose alone (2 mM), or in glucose (2 mM) plus lactate (5 mM) and pyruvate (0.1 mM). At the end of the experiment minimum and maximum NADH/NAD^+^ ratios were obtained by exposing the culture to 10 mM pyruvate and 5 mM sodium azide in the presence of 5 mM glucose and 10 mM lactate. Data are mean ± s.e.m. (3 cells), representative of 3 experiments.

In short, the cytosolic NADH pool in cultured neurons is impervious to moderate metabolic demand. This means that activated neurons adjust cytosolic NADH production to mitochondrial NADH demand in a seamless manner.

### Neuronal pyruvate and NADH in acute brain tissue slices

Next, we studied neurons in the adult somatosensory cortex. Firstly, control experiments were carried out in transgenic mice expressing GCaMP6 in neurons (Fig. 4A; Chen et al., 2013). Pipette-stimulation with an STB at about 50 - 100 µm of the imaged cells caused the expected tetradotoxin (TTX)-sensitive biphasic Ca^2+^ response (Fig. 4B). Sensitivity to TTX means that the Ca^2+^ signal is secondary to action potentials and not to local electroporation. The latter was confirmed independently by the observation that propidium iodide in the perfusate did not enter the recorded cells (data not shown). Pyruvate and NADH were assessed in brain tissue slices by expressing Pyronic and Peredox. To minimize confounding effects of tissue autofluorescence, we used Fluorescence Lifetime Imaging (FLIM) (Choi et al., 2012). As in cultured cells, the pyruvate pool of neurons in tissue slices was insensitive to electrical stimulation (Fig. 4C). At variance with the result in culture, there was no detectable change in pyruvate in response to extracellular lactate. This lack of response was not attributable to sensor saturation, as resting Pyronic occupancy in slices was 37 ± 16% (n = 3). A possible explanation for the insensitivity of pyruvate to extracellular lactate is that hypoxia in zones that are far from the slice surface can lead to higher resting ambient lactate (Mulkey et al., 2001). The NADH pool was not affected by the STB but did respond to tetanic stimulation (Fig 4D-E), suggesting that the balance between NADH production and consumption depends on the degree of activity and tissue context. As in cultures, exposure of slices to higher lactate increased neuronal NADH, although with slower kinetics: NADH remained stable for about a minute before slowly increasing. (Fig 4F). In order to investigate the source of NADH, lactate transport was inhibited with AR-C155858 (Díaz-García et al., 2017; San Martín et al., 2013). According to this protocol, a glycolytic origin should become evident as a larger NADH transient in the presence of the MCT blocker; conversely, a smaller transient would imply that the source of the NADH is extracellular lactate. However, the NADH transient evoked by tetanic stimulation was similar in the absence and presence of AR-C155858, with respective FLIM Δmean tau changes of 0.06 ± 0.04 (n = 20 slices) and 0.05 ± 0.03 (n = 5 slices; Figs. 4E and 4G). An alternative explanation for the activity-induced NADH transient could be the release of mitochondrial lactate, as discussed below.

**Figure 4.**
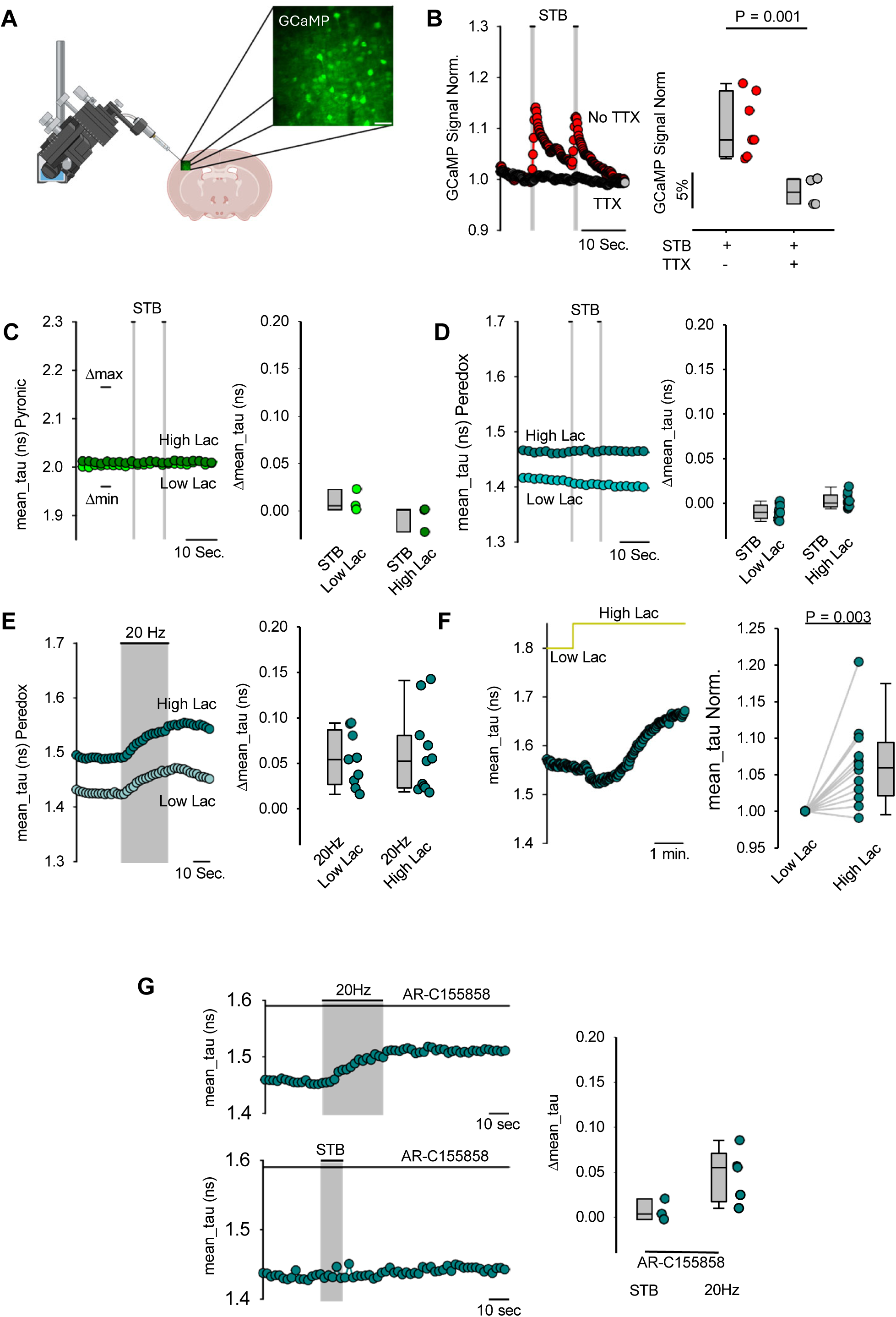
Neuronal pyruvate and NADH dynamics in brain tissue slices. **(A)** Neuronal activity was evoked by pipette stimulation at 50 -100 µm from the imaged cells. The inset shows a representative fluorescence two-photon image of neurons in the somatosensory cortex expressing the GCaMP sensor. Scale bar = 40 µm. **(B)** Effect of an STB on cytosolic Ca^2+^. Left panel: representative full-field GCaMP image capturing multiple neurons. Right panel: Summary of data in the absence (Red dots; n = 7 slices) and presence (gray dots; n = 4 slices) of tetrodotoxin (TTX; 1 µM). Slices were superfused with 2 mM glucose, 1 mM lactate, and 0.1 mM pyruvate. **(C)** Effect of an STB on cytosolic pyruvate in low (1 mM) and high (5 mM) extracellular lactate. Left panel: representative traces showing mean_tau of Pyronic. The minimum (min.) and maximum (max.) lines represent the mean_tau values obtained through a two-point calibration protocol used to establish the dynamic range of the sensor. Right panel: summary of data showing Δmean_tau (1 mM lactate, n = 3 slices; 5 mM lactate, n = 3 slices). The perfusion solution contained 2 mM glucose and 0.1 mM pyruvate. **(D)** Effect of an STB on cytosolic NADH in low (1 mM) and high (5 mM) extracellular lactate. Left panel: representative trace showing mean_tau of Peredox. Right panel: summary of data showing Δmean_tau (1 mM lactate n = 10 slices; 5 mM lactate n = 10 slices). The perfusion solution contained 2 mM glucose and 0.1 mM pyruvate. **(E)** Effect of tetanic stimulation (20 Hz for 30 s) on NADH in low (1 mM) and high (5 mM) extracellular lactate. Left panel: representative trace showing mean_tau of Peredox. Right panel: summary of data showing Δmean_tau (1 mM lactate, n = 9 slices; 5 mM lactate, n =11 slices). The perfusion solution contained 2 mM glucose and 0.1 mM pyruvate. **(F)** NADH response to switching from low (1 mM) to high (5 mM) extracellular lactate. Left panel: representative trace showing mean_tau of Peredox. Right panel: summary of data showing normalized mean_tau before and after lactate elevation (n = 12 slices). **(G)** Effect of electrical stimulation on NADH under MCT block. Left panels: representative traces showing mean_tau of Peredox in neurons exposed to tetanic stimulation (20 Hz for 30 s; top) or STB (bottom), both in the presence of AR-C155858 (1 μM). Right panel: summary data showing Δmean_tau in response to STB (n = 3 slices) and tetanic stimulation (5 = slices).

### Composition of pyruvate and NADH pools

Further to dissect the composition of the pyruvate and NADH pools, we simulated their behaviour using a mathematical model (Fig 5A; see equations in Methods). According to this model, the pyruvate pool is fed by glycolysis (Flux GLY), drained by mitochondrial consumption via the MPC (Flux MPC), and reversibly linked to lactate via LDH (K_1_) and to extracellular pyruvate via MCT (K_2_). The NADH pool is fed by glycolysis, drained by mitochondrial NADH uptake via MAS (K_4_), and sensitive to LDH activity (K_1_). The lactate pool is also connected to extracellular lactate (K_3_). In line with its equilibrium constant (K_eq_), the forward reaction of LDH (pyruvate to lactate) was set 14.2 x 10^4^ times faster than the reverse reaction (Williamson et al., 1967). MPC flux was assumed driven by demand, i.e. insensitive to increases in cytosolic pyruvate, and MAS flux was computed as a linear function of cytosolic NADH. The model is a system of four ordinary differential equations, five parameters and three external inputs. To solve the model we determined glucose consumption using a transport-block protocol (Bittner et al., 2010; San Martín et al., 2014; San Martín et al., 2022). In line with previous studies (Baeza-Lehnert et al., 2018; Sotelo-Hitschfeld et al., 2012) switching from low to high extracellular lactate (0.5 to 5 mM) reduced neuronal glucose consumption from 0.64 to 0.30 µM s^-1^ (Fig. 5B). Assuming negligible loss of carbons in glycolytic branches (Bolaños et al., 2008; Bouzier-Sore et al., 2006) these figures amount to 1.28 and 0.60 µM s^-1^ in respective pyruvate production. To convert Pyronic responses into cytosolic concentration, the FRET sensor was calibrated by exposing neurons to increasing concentrations of extracellular pyruvate, with lactate added in a 1:10 ratio to avoid loss of pyruvate through LDH. Following this procedure, pyruvate was estimated at 89 ± 4.2 μM in low lactate, and 266 ± 27 μM in high lactate. Using an analogous approach, resting concentration of lactate was determined at 880 ± 130 µM (Rauseo et al., 2026). Intracellular lactate was not quantified at high extracellular lactate conditions because available sensors have relatively low K_D_, precluding reliable measurements in high lactate concentration. By fitting the model to glycolytic pyruvate production, pyruvate concentrations and resting lactate concentration (Fig. 5D), the following results were obtained: Flux MPC= 1.18 µM s^-1^; K_1f_ = 9.3 x 10^4^ µM^-1^ s^-1^; K_2_ = 2.8 x 10^-4^ s^-1^; K_3_ = 2.8 x 10^-4^ s^-1^; K_4_ = 16.9 s^-1^. Thus, when neurons are in low lactate most glucose ends up metabolized in mitochondria and there is no contribution of lactate to the pyruvate pool. On the contrary, there is net export of monocarboxylates in the form of lactate, consistent with the effect of MCT1-2 blockage shown in Fig. 2B. Strikingly, the pyruvate pool grows substantially when cells are exposed to high lactate, even though glycolysis is inhibited. Consequently, lactate matches glucose as mitochondrial substrate (Figs. 5D-F). The impact of extracellular lactate on the NADH pool was identical, with lactate contribution rising to 0.62 µM s^-1^, that is, 51% of the total MAS flux (Fig. 5E).

**Figure 5.**
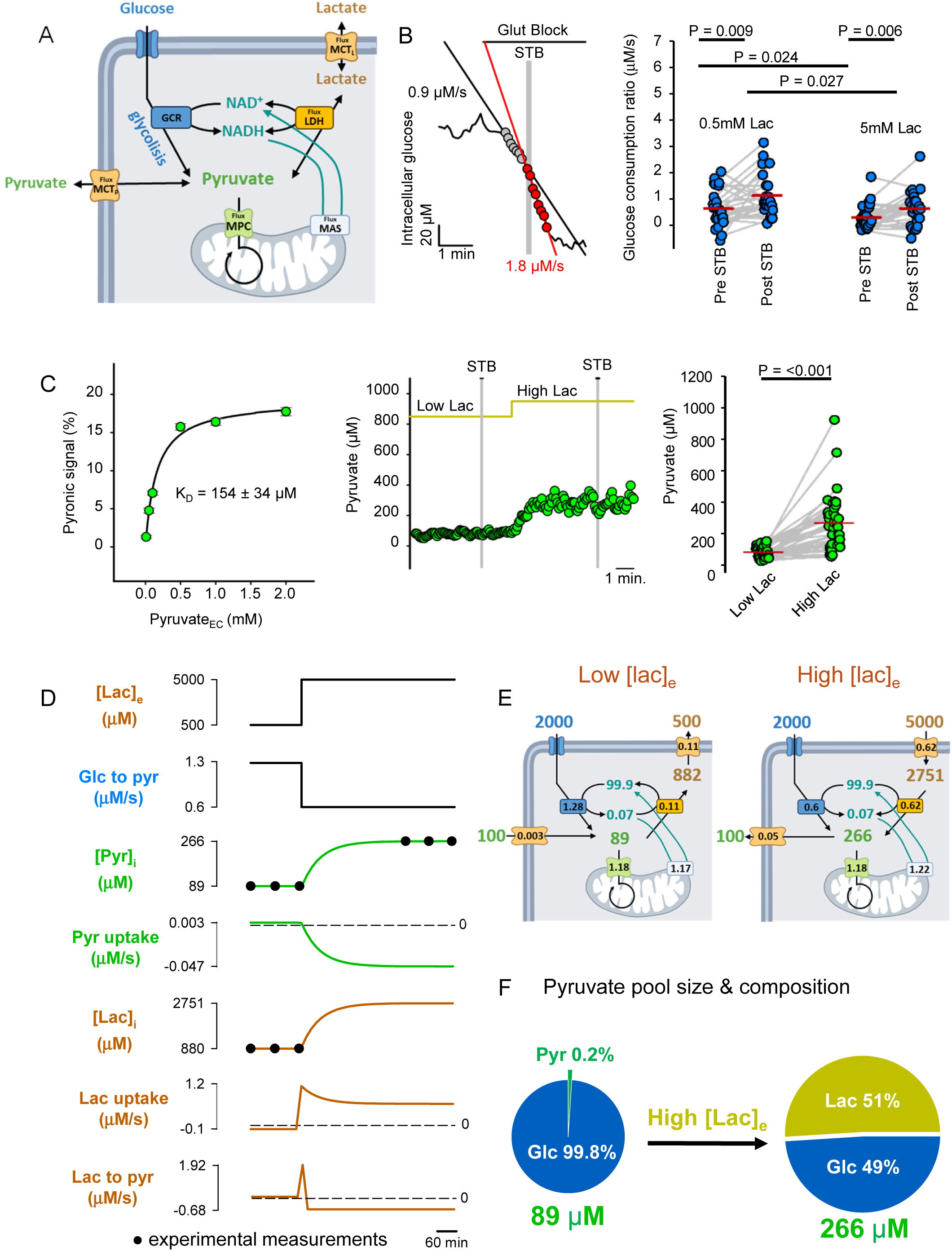
Simulation of neuronal fuelling: effect of extracellular lactate. Fuelling was solved by numerical simulation after experimental determination of glucose consumption and the steady-state concentrations of pyruvate and lactate. **(A)** Model of mitochondrial fuel dynamics. The cytosolic pools of pyruvate, NADH, and lactate are determined by the balance between glycolytic rate (GCR), mitochondrial pyruvate consumption (Flux MPC), NADH consumption by MAS, and the exchange of monocarboxylates at LDH and MCT. See differential equations in the Methods section. **(B)** The effect of an STB on the rate of glucose consumption was estimated in the presence of the GLUT blocker cytochalasin B (20 μM). Left panel: representative trace of a single neuron, recorded in the presence of 0.5 mM lactate. The STB is indicated with a gray bar. Right panel: summary of data from 27 cells in 4 experiments in low lactate (0.5mM) and from 28 cells in 4 experiments in high lactate (5mM). The superfusate contained 2 mM glucose and 0.1 mM pyruvate. **(C)** Quantitation of cytosolic pyruvate. Left panel: Pyronic calibration curve obtained by exposing neurons to increasing pyruvate concentrations (0.01 – 2 mM), each paired with a 10-fold higher lactate concentration (0.1 – 20 mM) to minimize LDH interference. Data from 30 cells across 3 independent experiments. Middle panel: representative calibrated trace from a single neuron STB-stimulated in the presence of low (0.5 mM) and high extracellular lactate (5 mM). Right panel: summary of 41 cells from 10 independent experiments. Cultures were superfused with 2 mM glucose and 0.1 mM pyruvate. **(D)** Dynamic response of neuronal metabolism to a step rise in extracellular lactate ([Lac]_e_), obtained by best fitting the model in Fig. 5A (equations in Methods) to the experimental data. Glycolytic pyruvate production (Glc to pyr) was inferred from measured glucose consumption rates; steady-state cytosolic pyruvate ([Pyr]_i_) and lactate ([Lac]_i_), were measured with Pyronic and Laconic (black dots). Predicted pyruvate and lactate uptake via plasma membrane MCTs, and lactate-to-pyruvate conversion at LDH are shown. **(E)** Steady-state solutions of neuronal metabolism at low and high extracellular lactate. Under low lactate, glucose is the near-exclusive fuel for mitochondria, with a small fraction being exported as lactate. Under high lactate, glycolysis is partially inhibited and lactate fluxes reverse. Fluxes are in μM·s⁻¹ and concentrations in μM. Arrows indicate flux direction. See model in Fig. 5A for reference. **(F)** Contribution of glucose and lactate to the cytosolic pyruvate pool. At low lactate (0.5 mM), the cytosolic pool of pyruvate amounts to 89 μM, which mostly originates from glycolysis. At high lactate (5 mM), the pyruvate pool swells to 266 μM, with similar contributions from glucose and lactate.

Next, we modelled the effect of electrical stimulation. The transport-stop protocol showed a 76% stimulation of glycolysis by the STB (Fig. 5B). Considering the change in glycolytic rate alongside unperturbed levels of pyruvate and NADH (from Figs. 2E and 3A) and lactate (Baeza-Lehnert et al., 2018), the rates of mitochondrial pyruvate and NADH uptake were predicted to increase by 83% (Fig. 6A). When the effect of STB was modelled at high extracellular lactate, the stimulations of pyruvate and NADH uptake were slightly weaker (58%; Fig. 6A). Conserved lactate influx in the face of augmented glycolytic flux resulted in a relatively lower contribution of lactate of 32%. These results are consistent with previous experimental data obtained at 1 mM lactate in the absence of pyruvate, in which the rate of neuronal glucose consumption was found to be stimulated to a similar extent by electrical stimulation, without detectable changes in the intracellular levels of pyruvate, lactate, glucose and ATP (Baeza-Lehnert et al., 2018).

In summary, the modelling of our experimental data indicate that extracellular lactate forces its way as mitochondrial fuel, by simultaneously inhibiting glycolysis while increasing the pyruvate and NADH pools, and that the substrate preference of neurons is insensitive to intrinsic neuronal activity and metabolic demand.

**Figure 6.**
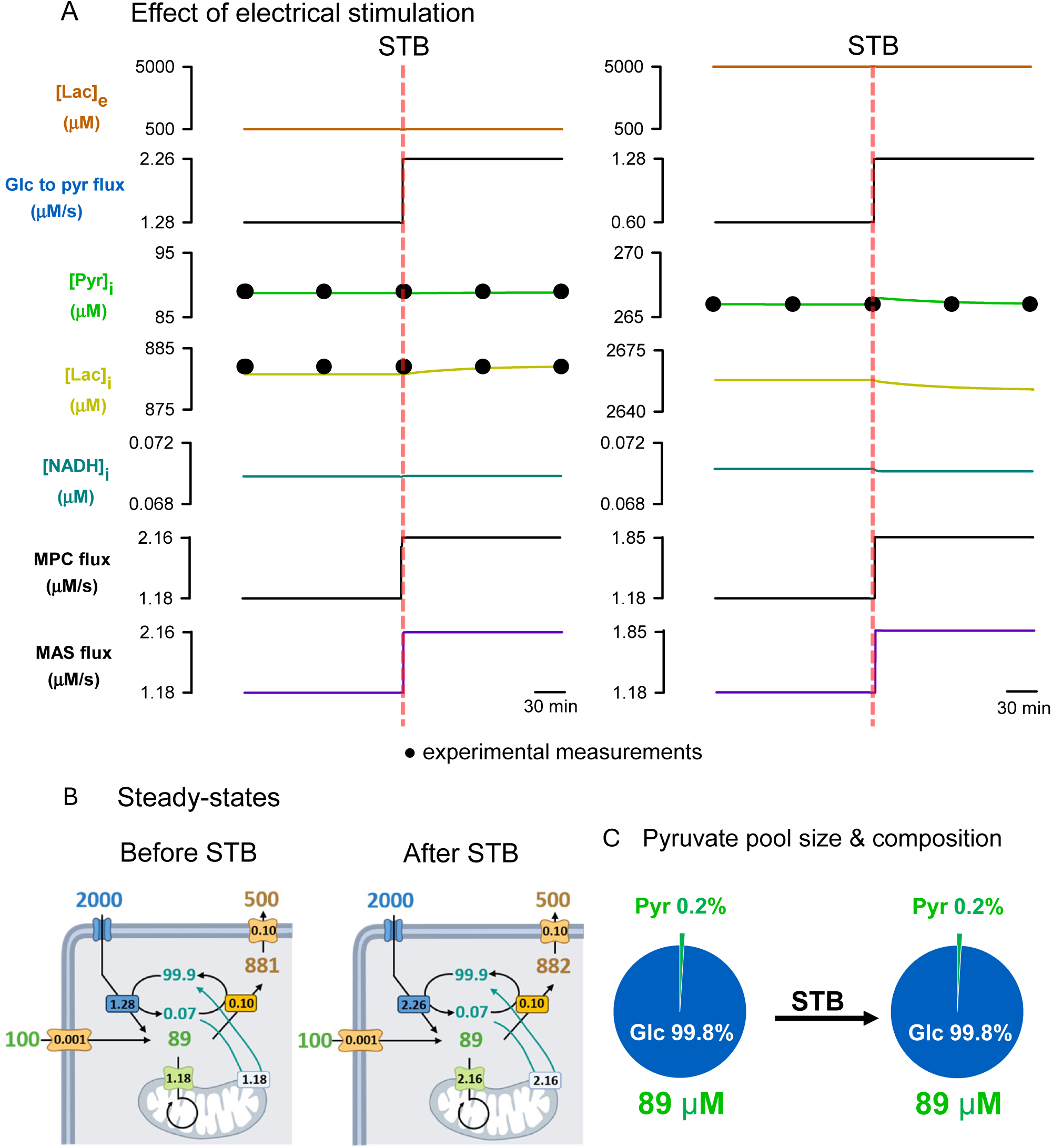
Simulation of neuronal fuelling: effect of electrical stimulation. Fuelling was solved by numerical simulation after experimental determination of glucose consumption and the steady-state concentrations of pyruvate and lactate. **(A)** Dynamic response of neuronal metabolism to an STB, obtained by best fitting the model in Fig. 5A (equations in Methods) to the experimental data, at low lactate (left panels) and high lactate (right panels). Glycolytic pyruvate production (Glc to pyr) was inferred from measured glucose consumption rates; steady-state cytosolic pyruvate ([Pyr]_i_) and lactate ([Lac]_i_), were measured with Pyronic and Laconic (black dots). Predicted NADH concentration, MPC flux and MAS flux are shown. **(B)** Steady-state solutions of neuronal metabolism before and after the STB. Electrical stimulation increases glucose consumption and mitochondrial pyruvate oxidation, while fluxes through MCTs and LDH remain unaffected. **(C)** No effect of the STB on the relative contribution of glucose and lactate to the cytosolic pyruvate pool. Electrical stimulation is without effect on the size of the cytosolic pool of pyruvate (89 μM), which is mostly originated from glycolysis.

## Discussion

Various cell types adapt to workloads through metabolic flexibility. For example, skeletal muscle cells and cardiomyocytes transition between fatty acids and glucose based on exercise intensity (Goodpaster & Sparks, 2017; Pascual & Coleman, 2016), while T lymphocytes shift heavily toward glucose upon antigen encounter (Wang & Green, 2012). Although the discovery of brain aerobic glycolysis suggested that neurons might similarly adjust fuel selection based on workload, our findings challenge this paradigm. Instead, we propose a nuanced, "opportunistic" model: neural activity dictates the overall metabolic pace, while the extracellular environment determines the fuel. By emphasizing the critical role of extracellular substrates over intrinsic neuronal properties, this insight reconciles long-standing, conflicting viewpoints on neurometabolic coupling (Bak & Walls, 2018; Barros & Weber, 2018).

### The neuronal fuelling debate

The adult brain is mostly energized by glucose, and neurons are its most abundant cell type. Therefore, it once seemed natural to assume that neurons are energized by glucose. However, the discoveries of activity-dependent local aerobic glycolysis in brain tissue (Fox et al., 1988; Prichard et al., 1991) and lactate production by stimulated astrocytes (Pellerin & Magistretti, 1994), led to the idea that active neurons may also be energized by lactate, formalized as the astrocyte-to-neuron lactate shuttle (ANLS) hypothesis. Consistently, high circulating lactate during exercise resulted in augmented lactate consumption and proportional reduction in glucose consumption by the brain (Quistorff et al., 2008). A fecund idea, ANLS has led to the discovery of new metabolic and signalling mechanisms in astrocytes, neurons and oligodendrocytes, in health and disease (Barros, 2013; Bonvento & Bolaños, 2021; Fernández-Moncada et al., 2024; Kuang et al., 2018; Looser et al., 2024; Magistretti and Allaman, 2018).

### Fuel selection is dictated by extracellular lactate and pyruvate, not by demand

Neurons may withstand a rise in energy demand and metabolic rate by up to 300% without detectable changes in cytosolic levels of ATP, ATP/ADP, glucose, pyruvate or lactate (Baeza-Lehnert et al., 2018; Rangaraju et al., 2014). This remarkable metabolic stability, which the present study extended to NADH, shows that neurons do not modify their metabolic profile in response to energy demand, they simply run at a faster rate. In stark contrast, fuel selection was found to be highly sensitive to external lactate availability. Supplying neurons with glucose, lactate, and pyruvate at physiological concentrations is relevant, since the standard practice of providing only glucose introduces a confounding factor by driving fast compensatory changes in redox balance, glycolytic flux and fuelling. Our data show that pyruvate and lactate pools are tightly linked by near-equilibrium LDH and MCT2, meaning that neurons operate close to the tipping point between monocarboxylate import and export and are ideally positioned to accommodate a rise in extracellular lactate.

### Where does extracellular lactate come from?

If fuelling preference is not an intrinsic property of neurons, focus shifts to their immediate milieu. Various sources of extracellular lactate are now recognized, local and systemic. Astrocytes and oligodendrocytes release lactate in response to neuronal cues, with K^+^ in charge of second-to-second regulation (Bittner et al., 2011; Looser et al., 2024), and glutamate, NO, NH ^+^, and adenosine involved in slower, longer-lasting effects (Barros et al., 2023; Bonvento & Bolaños, 2021; Theparambil et al., 2024). Hypoxia is a strong modulator of glycolysis and hypoxic pockets within cortical tissue may support physiological lactate production (Beinlich et al., 2024). Another source are mitochondria, which can release lactate under reducing conditions, a phenomenon exacerbated by hypoxia (Rauseo et al., 2026; Cárcamo-Lemus et al., 2025). Neurons and astrocytes have a similar density of mitochondria (Calì et al., 2019). It remains to be investigated whether the mitochondria of neurons and glial cells, and their processes, differ in their tendency to produce lactate. Neurons may also be reached by circulating lactate. At rest, both plasma and brain lactate are low and there is little exchange across the blood-brain barrier. However, during exercise plasma lactate increases and enters the brain to become a significant energy source, sparing glucose (Boumezbeur et al., 2010; Quistorff et al., 2008). The effectiveness of lactate was showcased in mice, where a severe hypoglycaemia was without functional consequence if lactate was infused in the circulation (Wyss et al., 2011).

### Limitations

Neurons in culture and brain slices lack vascular perfusion, systemic regulation, and complete network connectivity. Furthermore, our measurements were restricted to somata, which are metabolically distinct from axons and dendrites (Wei et al., 2023). Consequently, the core value of this study lies not in determining absolute concentrations and fluxes, but in highlighting the stark contrast between the metabolic impacts of energy demand and extracellular lactate and pyruvate availability.

## Acknowledgements

We thank Dr. Noemi Binini for technical support. We also acknowledge Chocolate, YC-B’s assistance dog, for his invaluable companionship and support. This work was supported by FONDECYT grants 1200029, 1230145 and 1260819, and Fondequip EQY240004.

## Materials and Methods

### Material

| REAGENT or RESOURCE | SOURCE/ IDENTIFIER |
| --- | --- |
| **Chemicals** |  |
| Sodium pyruvate | Sigma Aldrich, cat. #113-24-6/P2256 |
| Sodium L-lactate | Sigma Aldrich, cat #867-56-1/71718 |
| D-(+)-Glucose | Sigma Aldrich, cat. #G7021 |
| NaCl | Sigma Aldrich, CAS# 7647-14-5 |
| KCl | Sigma Aldrich, CAS# 7447-40-7 |
| CaCl<sub>2</sub> | Merck, CAS #100-04-8 |
| MgCl<sub>2</sub> | Merck, CAS #7791-18-6 |
| NaHCO<sub>3</sub> | Merck, CAS #144-55-8 |
| HEPES | Merck, CAS #7365-45-9 |
| Na<sub>2</sub>HPO<sub>4</sub> | Merck, CAS # 7558-79-4 |
| KH<sub>2</sub>PO<sub>4</sub> | Merck, CAS # 7778-77-0 |
| Phenol red | Merck, CAS # 143-74-8 |
| Sodium azide | Sigma Aldrich, CAS # 26628-22-8 |
| UK-5099 | MedChemExpress, CAS # 56396-35-1 |
| AR-C155858 | MedChemExpress, CAS # 496791-37-8 |
| Cytochalasin B | Millipore, CAS #14930-96-2 |
| TTX | Sigma-Aldrich, CAS #4368-28-9 |
| Pluronic™ F-127 | Molecular probes cat. #P6867 |
| Dimethyl sulfoxide | Sigma-Aldrich, CAS # D8418 |
| **Primary hippocampal cultures** |  |
| Isoflurane (100%, UPS grade) | Baxter cat. #10019036060 |
| Poly-L-lysine | Sigma Aldrich, cat. # P4832 |
| Trypsin-EDTA | Capricorn, Cat. #TRY-1B10 |
| Neurobasal-A medium | Gibco, cat. #A2477501 |
| D-Glucose | Sigma Aldrich, CAS #50-99-7 |
| Sodium pyruvate | Gibco, cat. #11360070 |
| Gluta MAX Supplement | Gibco, cat. #35050061 |
| B-27 Supplement | Gibco, cat. #17504044 |
| Penicillin/Streptomycin | Gibco, cat. #15140122 |
| **Genetically Encoded Sensors and Dye** |  |
| Pyronic | Addgene #51308 |
| Peredox | Addgene #32383 |
| CanlionicSF | Addgene #178342 |
| **FLII12Pglu-700uDelta6** | Addgene #17866 |
| **pAAV.Syn.NES.jRCaMP1a.WPRE.SV40** | Addgene #100848 |
| **Fluo4-AM ,cell permeant** | Invitrogen cat. #F14201 |
| **Virus generation** |  |
| **AAV helper-free vector systems** | Cell Biolabs. #VPK-400-DJ |
| **AAV purification kits** | Takara. Cat. #6666 |
| **HEK293FT cells** | Thermo Fisher Scientific, Cat# R70007 |
| **Polyethylenimine, Linear, (PEI 25K™)** | Polysciences, CAS #9002-98-6 |
| **Dulbecco's Modified Eagle Medium (DMEM) with high glucose** | Gibco #12100-038 |
| **Fetal bovine serum** | GE Healthcare. Cat. # SH103IH2540 |
| **Non-Essential Amino Acids (MEM-NEAA)** | Gibco, # 11140050 |
| **Geneticin™** | Roche, cat. # sc-29065A/G418 |
| **Software and Algorithms** |  |
| **Berkeley Madonna 8.3.18** | http://www.berkeleymadonna.com |
| **FluoviewFV10-ASW3.0** | Olympus |
| **Metafluor7.5.0.0** | Molecular Devices |
| **FLIMfit 5.1.1-19** | https://www.flimfit.org/ |
| **ImageJ1.54p** | http://imagej.net |

### Animals

All animal procedures were carried out in strict accordance with the recommendations in the Guide for the Care and Use of Laboratory Animals of the National Institutes of Health. Mice were housed at room temperature (20 ± 2 °C) in a 12 h light/12 h dark cycle with ad libitum access to food and water. Procedures performed in Valdivia were approved by the Centro de Estudios Científicos Animal Care and Use Committee (project 1230145) and sanctioned by the Dirección de Integridad, Seguridad y Ética de la Investigación, Universidad San Sebastián. Procedures performed in Zurich were approved by the local veterinary authorities and complied with the Swiss Animal Protection Law, Veterinary Office, Canton of Zurich (Animal Protection Act of 16 December 2005 and Animal Protection Ordinance 23 April 2008).

### Contact for reagent and resource sharing

Further information and requests for resources and reagents should be directed to the Lead Contact, L. Felipe Barros.

## Methods

### Primary hippocampal cultures

Primary hippocampal cultures were prepared from C57BL/6J × CBA/J mouse embryos. Pregnant mice at embryonic day 17.5 were euthanized by cervical dislocation following deep isoflurane anaesthesia, in accordance with institutional ethical guidelines and the ARRIVE principles. Under sterile conditions, 6-8 embryos were collected and immediately transferred to ice-cold HBSS supplemented with 5 mM glucose. Brains were isolated, and hippocampi were dissected under a stereomicroscope, taking care to completely remove the meninges.

Tissues was enzymatically dissociated in 1% trypsin-EDTA prepared in HBSS 1X (Hanks Balanced Salt Solution 10X: 100mM HEPES, 1.36 M NaCl, 74mM KCl, 50 mM Glucose, 44mM K_2_PO_4_, 50mg Phenol Red) for 15 minutes at 37°C. Digestion was stopped by adding Neurobasal medium supplemented with 10 mM glucose, 2% B-27 supplement, 1% GlutaMAX, and 5% fetal bovine serum (FBS). The tissue was then mechanically dissociated by gentle trituration. Dissociated hippocampal cells were plated on poly-D-lysine-coated glass coverslips (15 or 25 mm diameter) at a density of 160,000 cells/mL. Cells were allowed to adhere for 2 hours at 37°C in a humidified incubator (95% air, 5% CO₂). After attachment, the plating medium was replaced with serum-free Neurobasal medium supplemented with 10 mM glucose, 2% B-27, 1% GlutaMAX, and 10 μg/mL penicillin/streptomycin. Neuron-astrocyte co-cultures were maintained at 37°C in a humidified atmosphere (95% air, 5% CO₂), with two-thirds of the culture medium replaced every three days.

### Preparation of acute brain tissue slices

Mice were deeply anesthetized with isoflurane and euthanized by decapitation. Brains were rapidly extracted and placed in ice-cold, oxygenated cutting solution containing (in mM): 65 NaCl, 2.5 KCl, 0.5 CaCl₂, 7 MgCl₂, 25 glucose, 105 sucrose, 25 NaHCO₃, and 2.5 Na₂HPO₄. Coronal brain slices (300 µm thick) were prepared using a Microm HM 650 V vibratome. Slices were immediately transferred to oxygenated artificial cerebrospinal fluid (aCSF) composed of (in mM): 126 NaCl, 3 KCl, 2 CaCl₂, 2 MgCl₂, 25 glucose, 26 NaHCO₃, and 1.25 NaH₂PO₄, maintained at 34 °C. Slices were allowed to recover for 1 hour before being used for imaging experiments. All solutions were continuously oxygenated with a gas mixture of 95% O₂ and 5% CO₂ and were freshly prepared on the day of the experiment.

### Expression of fluorescent sensors

Adeno-associated viral vectors (AAVs) were utilized to express genetically encoded fluorescent sensors in cultured cells and in vivo. Viral particles used in culture were produced in-house using the AAVpro Helper Free-vector System and purified using a modified protocol based on the AAV purification kits from Takara Bio. Neuronal specificity was achieved with the human synapsin promoter (hSyn). Viral particles were added to the culture medium 2-5 days before imaging, obtaining sensor expression in >80% of neurons.

For brain tissue expression stereotaxic surgery was performed in female wild-type mice (C57BL/6J; Charles River) of 8–10 weeks of age (20- 25g body weight). Two 1 × 1 mm craniotomies were created above the somatosensory cortex using a Bien-Air Dental dental drill. 50 µL of adeno-associated virus (AAV) vectors containing the respective GEFIs (titer 1.02 × 10^11 VG/ml; Viral Vector Core Facility [VVF], University of Zürich) were injected into the somatosensory cortex at a cortical depth of 300 µm. Care was taken to avoid large vessels to prevent bleeding. In addition, a transgenic GCaMP mouse with a neuronal targeting signal was used. This mouse was provided by Dr. Aiman Saab from the Institute of Pharmacology and Toxicology, University of Zurich.

### Fluorescent dye loading

Cultures were incubated for 15 min at 37 °C under 95% O₂/5% CO₂ with the calcium indicator Fluo-4 AM (4 µM; 1:4000) prepared in KRH buffer supplemented with 0.02% Pluronic F-127, 2 mM glucose, 0.5 mM lactate and 0.1 mM pyruvate. Cultures were then washed for 15 min at room temperature in the same buffer lacking Pluronic F-127 before imaging.

### Fluorescence microscopy

Cultures were imaged at either room temperature (22–25 °C) or 34–35 °C in a solution equilibrated with 95% air/5% CO₂ and containing (in mM): 112 NaCl, 3 KCl, 1.25 CaCl₂, 1.25 MgSO₄, 1–2 glucose, 10 HEPES, and 24 NaHCO₃ (pH 7.4). Confocal imaging was performed using an Olympus BX61WI FV1000 upright confocal microscope equipped with a 20x water-immersion objective (NA 0.95) and 440, 488, and 543 nm laser lines. Fluorescence imaging was also performed using an Olympus BX51 upright microscope equipped with a CAIRN monochromator and a 20x water immersion objective (NA 1.0; Faversham, UK) and a Rollera camera controlled by MetaFluor software.

Tissue slices were imaged using a custom-built two-photon laser scanning microscope (2PLSM) (Mayrhofer et al., 2015) with a tunable pulsed laser (MaiTai HP system, Spectra-Physics and Chameleon Discovery TPC, Coherent Inc) and either a 20× water-immersion objective. (W Plan-Apochromat 20 x/1.0 NA, Zeiss) or 25× (W Plan-Apochromat 25 x/1.05 NA, Olympus)

### Sensor and fluorescent dye imaging

FRET sensors (Pyronic and FLII¹²Pglu-700µΔ6) were imaged using 440 nm excitation, and emission was collected at 480/15 nm and 550/15 nm. The ratiometric NADH/NAD⁺ sensor Peredox, based on T-Sapphire and mCherry, was imaged using a monochromator with 405/10 nm excitation for T-Sapphire and 575/10 nm excitation for mCherry, with emission collected using 524/25 nm and 628/25 nm band-pass filters, respectively. Single-fluorophore sensor (CanlonicSF) and the Ca²⁺ dye (Fluo-4 AM) were imaged using 488 nm excitation and 515 ± 10 nm emission.

FRET data are presented as fluorescence ratios (mTFP/Venus for the pyruvate sensor and YFP/CFP for the glucose sensor). The ratiometric Peredox signal was calculated as T-Sapphire/mCherry. And signals or fluorescence ratios were normalized to the minimum intensity value. All experiments were initiated from a steady-state baseline established with a reference buffer containing 2 mM glucose, 0.5 mM lactate, and 0.1 mM pyruvate.

### Electrical Field Stimulation

Mixed cultures of hippocampal neurons and astrocytes grown on 15 mm coverslips were field-stimulated using a RC-21BRFS chamber (Warner Instruments) and a PRO-4 device (World Precision Instruments, WPI). Pulses (50mA output) were generated with a WPI A385 High Current Stimulus Isolator connected to a WPI A382 Battery Charger. The stimulation protocol was a short theta burst that mimic hippocampal activity, consisting of two trains of impulses separated by 10 seconds. Each train lasted for 1 second and was composed of twenty pulses of 1 ms duration, distributed into five groups of four at 0.2-second intervals, with each pulse spaced 0.01 seconds apart (illustrated in Fig. 3A). Short theta burst (STB) stimulation elicited a characteristic biphasic Ca²⁺ transient (Fig. 3B), followed by a slower increase in neuronal Na⁺ levels, which has been shown to correlate with the bulk of ATP demand (Baeza-Lehnert et al., 2018). Alternatively, the cultures were exposed to increasing stimulation in terms of duration and intensity: 1 pulse, 5 pulses, 10 pulses, 20 pulses, 40 pulses (STB), or 600 pulses (20Hz for 30 s).

### Local Electrical Stimulation and Temperature Control

Local stimulation in acute slices (Fig. 3D) was delivered using a Multi-Channel Systems STG3008-FA stimulator connected to a Millar Instruments catheter (EPR-800/801/802 series), positioned 50-100 µm from target cells via a motorized micromanipulator. The system was integrated with an NPI Electronic TC-20 temperature controller (19" rack-mounted, dual independent channels) maintaining 37°C ± 0.1°C through integrated Peltier elements, with continuous monitoring via a bath probe. The sensitivity to tetrodotoxin (TTX) shows that the Ca2+ transient is secondary to action potential and not to local electroporation (Fig 3E). The latter was confirmed independently by the observation that the extracellular dye did not enter the recorded cells (data not shown).

Further to investigate the metabolic response of neurons to neurotransmission under physiological conditions, we studied neurons in acute brain slices from the adult somatosensory cortex. This preparation preserves the structural and metabolic complexity of the brain, allowing us to assess neuronal responses in a more physiologically relevant context. As a control, we first validated the neuronal response to electrical stimulation in transgenic mice expressing GCaMP6, a genetically encoded calcium indicator. Pipette stimulation with a STB protocol at a distance of 50–100 µm from the imaged cells elicited the expected TTX-sensitive biphasic Ca²⁺ transient (Fig. 3D-B). TTX is a potent voltage-gated sodium channel blocker, which prevents action potential generation. The sensitivity of the Ca²⁺ response to TTX confirmed that neuronal activation was mediated by action potentials rather than direct electroporation. This was further supported by the absence of extracellular dye uptake in stimulated cells (data not shown), ruling out membrane disruption. The robust Ca²⁺ transients observed in these experiments indicated that neuronal activation was successfully achieved using the STB protocol.

### Pyronic calibration

To determine Pyronic K_D_, the sensor was calibrated on the cytosol of neurons. To minimize the influence of LDH activity, an LDH clamp protocol was used in which pyruvate and lactate were applied at a fixed 1:10 ratio, maintaining LDH close to equilibrium (Glancy et al., 2021). Neurons were perfused with increasing pyruvate/lactate concentrations while maintaining this ratio (0.01:0.1; 0.05:0.5; 0.1:1; 0.5:5; 1:10; 2:20 mM). LDH equilibrium during calibration was monitored using the NADH/NAD⁺ sensor Peredox.

### Transport-stop assays for flux measurements

Fluxes were measured using genetically encoded FRET sensors following acute pharmacological disruption of metabolite steady states with transport blockers, as previously reported. (Bittner et al., 2010; Barros et al., 2013). Glucose consumption rate was computed in the presence of the GLUT blocker cytochalasin B (20 µM) before and after STB stimulation. Mitochondrial pyruvate consumption was measured in the presence of the MPC blocker UK-5099 (1 µM) before and after STB stimulation. Linear regressions were used to estimate fluxes. The onset of flux stimulation was computed as the intersection between linear regressions before and after STB.

### Mathematical modelling

To simulate cytosolic metabolite dynamics a mathematical model was implemented in Berkeley Madonna (version 8.3.18). The model consists of a system of four coupled ordinary differential equations, which were numerically solved using the Rosenbrock integration method:

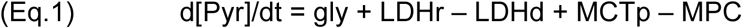

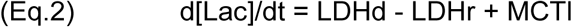

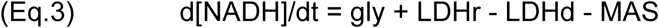

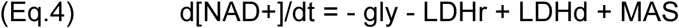

Where LDHd = K1f x Pyr x NADH; LDHr = K1r x Lac x NAD+; MCTp = K2 x (Pyre - Pyrin); MCTl = K3 x (Lace – Lacin); MAS = K4 x NADH; Pyr = Pyruvate and Lac = Lactate. Gly represents the glycolytic flux beyond GAPDH (glucose consumption x2). Lactate dehydrogenase fluxes were separated into the reductive direction (LDHr) and oxidative direction (LDHd). The reverse LDH rate constant was defined as K1r = K1f x 0.0000704; The equilibrium constant (0.00011) was taken from Williamson et al., (1967) at 38°C and pH 7.0, and adjusted to pH 7.2 (corresponding to 64 nM [H⁺]), yielding 0.00011 × 0.64 = 0.0000704. Monocarboxylate transporter fluxes (MCTl and MCTp) describe transmembrane exchange of pyruvate and lactate. MPC represents mitochondrial pyruvate uptak. MAS represents the shuttling of NADH electrons.

### Statistical Analysis

Data are expressed as mean ± SEM. Data analysis was performed in Sigma Plot software. A paired t-test was applied to ascertain differences in before-after treatment protocols with normal distribution. In case of failed normality test (Shapiro-Wilk), differences were assigned with the Mann Whitney-Wilcoxon signed rank test (pairs). *, p < 0.05. Not significant (NS), p > 0.05).

